# Effect of quercetin and kaempferol isolated from the *Ginkgo biloba* L. suspension culture extract on lipofuscin accumulation

**DOI:** 10.1101/2023.05.10.540159

**Authors:** Irina Sergeyevna Milentyeva, Anastasia Mikhailovna Fedorova, Anna Dmitrievna Vesnina, Alexander Yurievich Prosekov, Ludmila Konstantinovna Asyakina, Olga Alexandrovna Neverova

## Abstract

Lipofuscin, known as an age pigment, is an autofluorescent lipopigment formed by lipids, metals, and misfolded proteins that accumulates in nerve cells, heart muscle cells, and skin. Individual bioactive compounds obtained from medicinal plant extracts reduce the level of lipofuscin in *C. elegans* nematodes in *in vivo* experiments. We aimed to study the effect of quercetin and kaempferol isolated from the *in vitro* suspension culture extracts of *Ginkgo biloba* L. on the accumulation of lipofuscin. According to the results, 200 μM concentrations of these bioactive compounds showed the maximum decrease in lipofuscin levels in *C. elegans* N2 Bristol during their 15-day cultivation, compared to the control. Thus, our study showed that quercetin and kaempferol are able to reduce this age pigment in *in vivo* experiments.

## Introduction

Aging is a process by which the viability of an organism decreases under the influence of various mechanisms at all levels (molecules, tissues, etc.), with weakening adaptation and a growing likelihood of disease [1].

The accumulation of lipofuscin is the main factor that limits the cells’ functionality during aging and reduces their lifespan. In the human body, lipofuscin accumulates mainly in resting or postmitotic cells: cardiomycetes, retinal cells, neurons, skeletal muscles, and beta cells [2]. Lipofuscin is considered as intralysosomal “garbage” or waste material that comprises 30-58% of proteins, 20-50% of lipo-like substances, and up to 20% of trace metals, including Fe^3+^, Fe^2+^, Cu^2+^, Zn^2+^, Al^3+^, Mn^2+^, and Ca^2+^ [3].

Lipofuscin produces various biological effects on the body. It is known to inhibit the proteasome system, which is associated with the activation of *HDAC/HO1* or the transcription factor *AP1*. This inhibitory effect decreases as free peptide ends are removed [4]. The chemical activity of lipofuscin lies in its ability to increase in size due to reactive groups located on the surface. As a result, lipofuscin can react with other proteins, particularly with ubiquitin, a protein that can modify the functions of other proteins [5].

As mentioned earlier, lipofuscin is a hallmark of cellular aging. Its distribution in the brain of normal-aged mammals defines a specific aging pattern that correlates with altered neuronal cytoskeleton and cell movement. With age, the adult brain becomes overloaded with intraneuronal deposits of lipofuscin and neuromelanin pigments [3].

However, the number of lipofuscin aggregates increases not only with age, but also in diseases associated with neurodegenerative disorders (neuronal ceroid lipofuscinosis, age-related molecular degeneration, Parkinson’s, Alzheimer’s, and Huntington’s diseases, etc.). Neuronal ceroid lipofuscinosis, which results from the deposition of “ceroid lipofuscin”, is associated with mutations in 14 genes (from CLN1 to CLN14) and manifests itself through lipid abnormalities, inflammation, and lysosomal deposits of lipofuscin aggregates [6]. The accumulation of lipofuscin also leads to age-related molecular degeneration, i.e., a degenerative disorder of the pigment epithelium of the central retina, which is the cause of vision loss in the elderly [7]. In Parkinson’s disease, lipofuscin, together with α-synuclein, contributes to the degeneration of dopaminergic neurons. Neurons that contain lipofuscin are more susceptible to pathologies, such as changes in the cytoskeleton (lower levels of cytostructural non-phosphorylated protein of neurofilaments) and the deposition of α-synuclein [8]. The early stages of Alzheimer’s disease are characterized by damaged pyramidal neurons that accumulate lysosomes in the hippocampus and prefrontal cortex. The activation of autophagocytosis causes lysosomes to acquire aggregates of lipofuscin. When neurons in the extracellular space were destroyed, aggregates of lipofuscin were found with β-amyloid plaques. Presumably, lipofuscin is responsible for aluminum binding [9]. The brain of people suffering from Huntington’s disease contains increased amounts of extraneuronal deposits of lipofuscin in pyramidal neurons [10].

The nematode worm *Caenorhabditis elegans* is a universal model system used in biomedical and toxicological studies. *C. elegans* is an ideal model for aging research, since it has a short lifespan (2–3 weeks) and fast reproducibility (3–4 days). In addition, its signaling pathways are similar to those in the human body. For example, the *daf-2* gene, one of the most studied genes in *C. elegans’s* genome, gave rise to the genes for insulin and the receptor for insulin-like growth factor-1 in humans [11]. Lipofuscin is an autofluorescent material that is used as a marker of cellular and organismal aging, particularly in *C. elegans*. When excited by UV or blue light, lipofuscin fluoresces in the wavelength range from yellow to red. In *C. elegans*, lipofuscin accumulates with age and also with oxidative damage. The “blue” (350/460 nm) autofluorescence does not increase much age, peaking only before the death of the organism. The “red” (546/600 nm) autofluorescence increases linearly with time and correlates with life expectancy. The “green” (470/525 nm) autofluorescence combines the characteristics of the “red” and the “blue” ones [12]. There are studies that aim to discover new pharmacological agents that reduce the accumulation of lipofuscin. Plant extracts containing bioactive substances (flavonoids, phenolic acids, ginsenosides, essential oils, etc.), as well as individual substances, have been found to effectively reduce the level of lipofuscin in the nematodes *C. elegans* N2 Bristol in *in vivo* experiments. For example, 50 μg/ml of birch extract (*Betula utilis*) [13], 1000 μg/ml of garden purslane extract (*Portulaca oleracea* L.) [14], and 80 mg/ml of raspberry extract [15] reduced lipofuscin by 16.5–47.1% in nematodes. Wang *et al*. [16] reviewed a study where peptides from sesame oil cake at a concentration of 12.5 μg/ml reduced the accumulation of lipofuscin by 71.5%. Another study showed a similar effect of 180 μM of rosmarinic acid [17].

Although most bioactive substances and extracts reduce the level of lipofuscin in nematodes and increase their lifespan, not all polyphenolic compounds have been studied for the accumulation of this age pigment. These compounds include quercetin and kaempferol. Quercetin (C15H10O7) is classified as a flavonol, one of six subclasses of flavonoid compounds. Quercetin is found in a variety of foods, including apples, berries, Brassica vegetables, capers, grapes, onions, tea, and tomatoes. Quercetin is also found in medicinal plants such as *Ginkgo biloba* L., *Hypericum perforatum* L., and *Sambucus canadensis* L. [18]. Li *et al*. [19] reported that quercetin has anti-inflammatory properties and an immunosuppressive effect on dendritic cell function. It also lowers blood pressure and exhibits antioxidant, antiviral, antitumorous, antimicrobial, neuroprotective, and cardioprotective effects [20]. Kaempferol (C15H10O6), also classified as a flavonol, is found in many products (capers, saffron, fodder cabbage, brown mustard, etc.) and medicinal plants (*Ginkgo biloba* L., *Aloe Vera* L. EX WEBB, *Pulmonaria officinalis* L., *Panax ginseng* C.A. Mey, etc.) [21]. Most well-known are its anti-inflammatory and antitumorous effects [22]. In addition, kaempferol has antibacterial, antioxidant, neuroprotective, cardioprotective, and antidiabetic effects [23].

We aimed to study the effect of quercetin and kaempferol isolated from the extract of the suspension culture of *Ginkgo biloba* L. on the accumulation of lipofuscin.

## Study objects and methods

Quercetin and kaempferol were isolated from the extract of the suspension culture of *Ginkgo biloba* L. The earlier stages of research involved obtaining a suspension culture of *Ginkgo biloba* L. and its aqueous-alcoholic extract, followed by the isolation of individual bioactive substances from the extract. First, a callus culture of *Ginkgo biloba* L. was obtained from its sterile seeds as described by Le *et al*. [18]. The primary callus was separated from the remnants of plant explants, transferred to a fresh nutrient medium and cultivated for 4–5 weeks. To obtain cell suspensions, 300–400 mg amounts of *Ginkgo biloba* L. callus culture were placed in 250 ml flasks with 25 ml of liquid nutrient medium and cultivated on a shaker-incubator (LSI-3016A) at 95–100 rpm. After 18–20 days of cultivation, the supernatant fraction of cells was reseeded by gradually increasing the dilution from 1/2 to 1/8 (the inoculum-to-medium ratio) and shortening the subcultivation cycle. Suspension cultures were grown in 250 ml flasks (30–40 ml of suspension, 100 rpm) on the MS (Murashige-Skoog) medium in combination with growth regulators 2,4-D (2.00 mg/l) and kinetin (0.10 mg/l). The growth cycle was 2–3 weeks. All the cell cultures under study were cultivated in a climate chamber (Binder, Germany) under sterile conditions, in the dark or in the light (16-hour daylight), on a stationary circular shaker in the dark at 26–27 °C, 70–75% humidity, and 95–100 rpm.

The aqueous-alcoholic extract of the suspension culture of *Ginkgo biloba* L. [24] was obtained, and bioactive substances (quercetin and kaempferol) were isolated from it, as described by Le *et al*. [18]. The extraction was performed for 6 hours at 55 °C, with 70% ethanol. Quercetin and kaempferol were isolated by high performance liquid chromatography (HPLC) on a liquid chromatograph (Shimadzu LC-20 Prominence, Japan) and purified (95%) (Figure 1).

**Figure 1.**
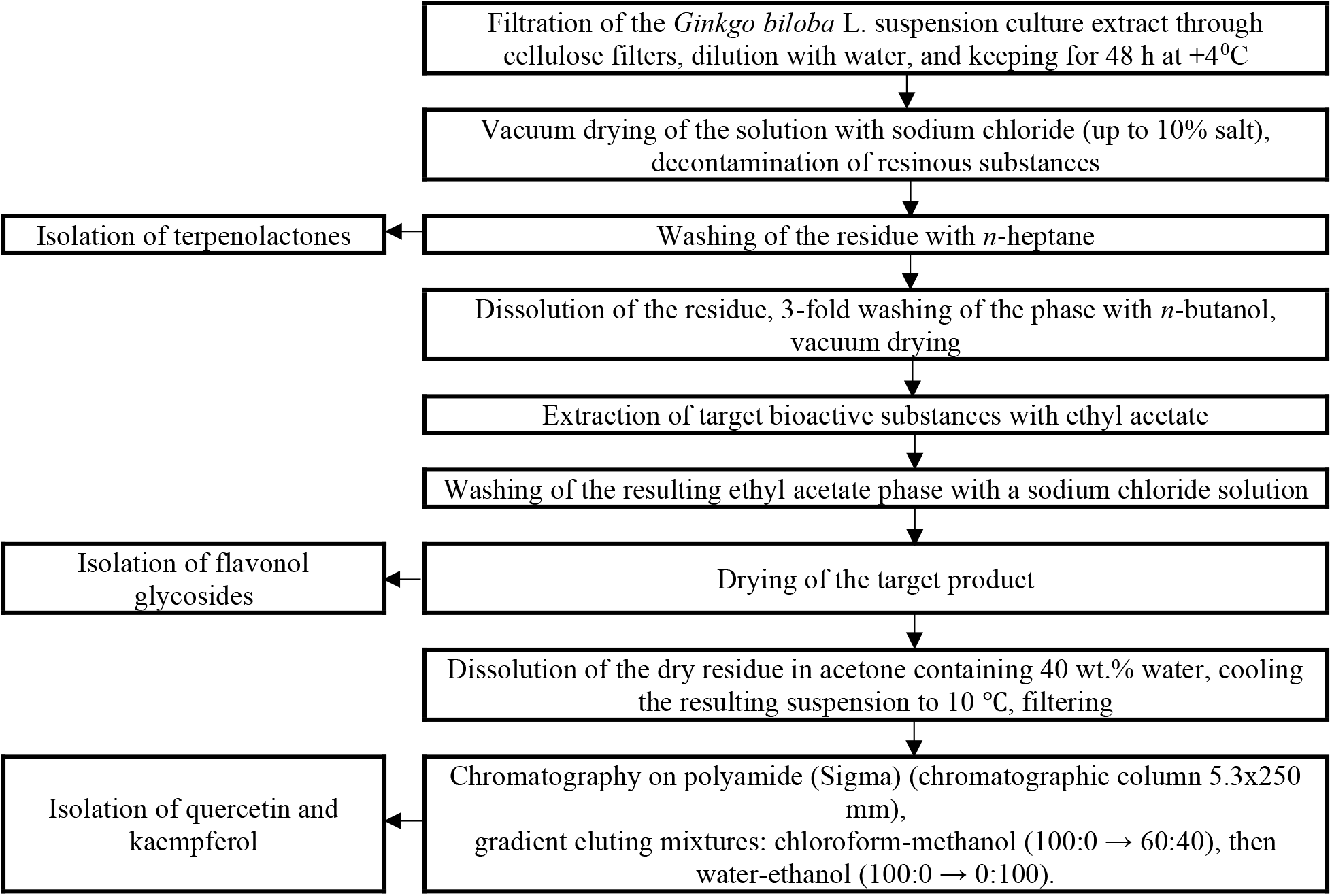
Isolation and purification of quercetin and kaempferol from the extract of the *Ginkgo biloba* L. suspension culture

**Figure 1.**
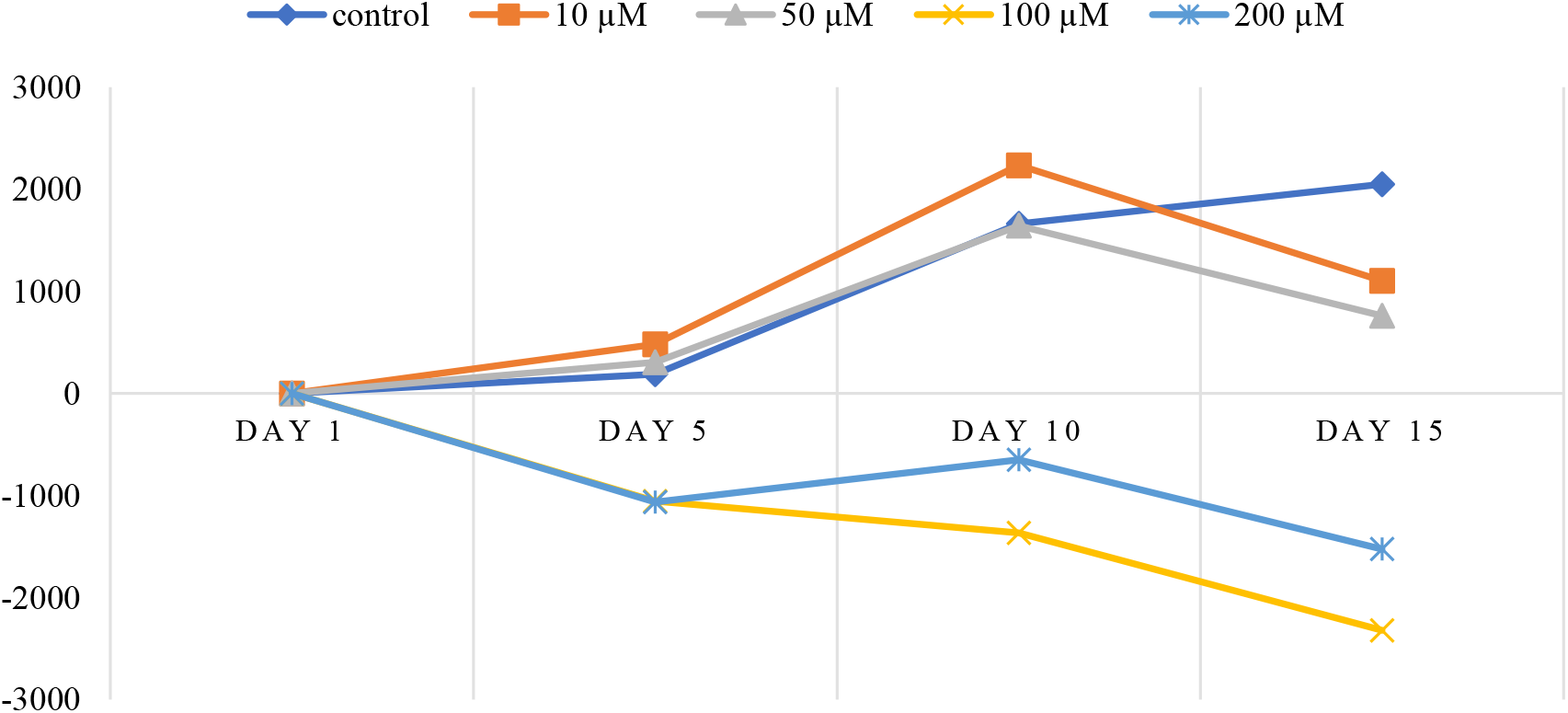
Accumulation of lipofuscin in *C. elegans* treated with quercetin isolated from the *Ginkgo biloba* L. suspension culture extract

The effect of quercetin and kaempferol on lipofuscin accumulation was studied by using the nematode *Caenorhabditis elegans* of the wild strain N2 Bristol as a model organism. The nematodes were provided by the Laboratory for the Development of Innovative Medicines and Agrobiotechnologies, Moscow Institute of Physics and Technology (National Research University) (Dolgoprudny, Russia).

*Escherichia coli* OP-50 bacteria (V.A. Engelhardt Institute of Molecular Biology, Russian Academy of Sciences, Moscow, Russia) were used as a nutrient for nematodes. The bacteria were added once, at the beginning of the experiment, to a final concentration of 5 mg/mL.

Nematodes were cultivated on solid NGM agar (1000 ml ddH2O, 2.5 g bactopeptone, 3 g NaCl, and 17 g bactoagar autoclaved at 120 °C for 15 min and then mixed with 1 ml 1M CaCl2, 1 ml 1M MgSO4, 1 ml 5 mg/ml cholesterol, 25 ml K2HPO4 + KH2PO4, pH 6.0). For this, 50 μl of *E. coli* OP50 overnight culture was placed in the center of a 100 mm Petri dish under sterile conditions. Then, a drop of the bacterial culture was rubbed with a sterile glass rod around the center of the dish in the shape of a square, without touching the walls. The dishes were incubated for 24 hours at 37 °C in a climate chamber (Binder, Germany). Then, the nematodes were transferred onto a bacterial lawn on new NGM agar plates under sterile conditions using a nematode transfer loop that was preliminarily annealed on fire and cooled. Then, a 0.5 cm x 0.5 cm piece of agar with nematodes was cut out with a sterile scalpel, transferred into the center of a new plate top-down, and incubated at 20°C.

Next, the nematodes were synchronized. For this, 5–10 ml of sterile water was added to an agar Petri dish with nematodes and pipetted several times until all nematodes and nematode eggs were no longer adhesively attached to the agar. The liquid was transferred from the Petri dish into a 50 ml tube and centrifuged for 2 min at 1200 rpm (280 g). The supernatant was removed and the precipitate was washed with 10 ml of distilled water, after which the process was repeated. The supernatant was removed and 5 ml of a freshly prepared mixture (1 ml of 10 N NaOH, 2.5 ml sodium hypochlorite, and 6.5 ml water) was added to the precipitate. It was well mixed and intensively vortexed for 5 minutes with a break every 2 minutes, while the hydrolysis of nematodes was observed under a microscope. At the end of the process, 5 ml of M9 medium (1000 ml of ddH2O, 3 g of KH2PO4, 5 g of NaCl, 6 g of Na2HPO4, and 1 ml of 1 M MgSO4) was quickly added to neutralize the reaction. After centrifugation for 2 minutes at 2500 rpm (1100 g), the supernatant was removed and the precipitate was resuspended in 10 ml of fresh sterile water, followed by three repetitions of neutralization and centrifugation. The fourth precipitate was washed with 10 ml of an S-medium (1000 ml S base, 10 ml 1 M potassium citrate, pH 6, 10 ml trace metal solution, 3 ml 1 M CaCl2, 3 ml 1 M MgSO4). The supernatant was removed and 10 ml of the S-medium was added. The test tube with nematode eggs was left on a shaker for 24 hours at room temperature until the nematodes entered the L1 development stage.

Then, the nematodes were cultivated in a liquid medium. For this, we added an overnight bacterial culture of *E. coli* OP50, which was previously washed from LB-broth (1000 ml ddH2O, 5 g NaCl, 10 g tryptone, 5 g yeast extract) and resuspended in the S-medium to a final bacterium concentration of 0.5 mg/ml. Then, 120 μl of the suspension of the nematodes with the bacteria was introduced into each well of a 96-well plate (Eppendorf, USA). The plate was sealed with a film and left for 48 hours for incubation in a climatic chamber (Binder, Germany) at 20 °C. After 48 hours, 15 μl of 1.2 mM 5-fluoro-2-deoxyuredin (FUDR) was added into each well to prevent the worms from reproducing. The plate was left for 24 hours at 20 °C until the worms reached the L4 stage. Then, 15 μl amounts of the tested compounds were added to the nematodes in line with the experimental plan.

The bioactive substances isolated from the *Ginkgo biloba* L. suspension culture extract were prepared as follows. First, quercetin and kaempferol stocks were prepared in 10 mmol/l or 50 mmol/l of dimethyl sulfoxide (DMSO). Then, each substance was titrated by diluting the stock solution with distilled sterile water to the stock concentrations of 200 μM, 100 μM, 50 μM, and 10 μM.

The accumulation of lipofuscin in *C. elegans* N2 Bristol was measured by spectrofluorimetric analysis. For this, 135 μl amounts of the L1 synchronized nematodes in the S-medium with 5 mg/ml of *E. coli* OP-50 were dropped into the wells of a 96-well plate in 6 repetitions for each tested concentration of the bioactive substances. The plates were incubated at 25 °C for 48 hours until the nematodes reached the L4 stage. Then, 15 μl of FUDR was added to each well and the plates were incubated for another 24 hours at 25°C. After that, 15 μl amounts of the tested substances were added to the wells at concentrations of 10, 50, 100, and 200 μM. The plates were incubated in a climatic chamber (Binder, Germany) at 25°C for 24 hours, after which the first cycle of spectrometric measurements was performed to assess the effect of the tested substances on lipofuscin accumulation in the nematodes.

The measurements were carried out on a Clariostar spectrophotometer (BMG Labtech, Germany) with the following settings: measurement type – fluorescence intensity (FI); microplate type – 96-well plates; input wavelengths of excitation and radiation – 340 and 420 nm, respectively; autofluorescence reading direction – bidirectional, horizontal from left to right, top to bottom. For the measurements, we placed the experimental 96-well plate with nematodes in the spectrophotometer, adjusted the resolution and focal height, checked the program settings, and started the process. This procedure was repeated to measure autofluorescence in the subsequent plates. The results were automatically exported to the Excel program, which was used to process the data.

## Results and discussion

Table 1 shows the average values of lipofuscin autofluorescence at 340/420 nm in *C. elegans* after 1, 5, 10, and 15 days of incubation following their treatment with quercetin and kaempferol isolated from the *Ginkgo biloba* L. suspension culture at concentrations of 10, 50, 100, and 200 μM in six repetitions, as well as standard deviations.

**Table 1.** Autofluorescence of lipofuscin in the nematodes treated with bioactive substances

| Nematode incubation time, days | Concentration of bioactive substance, $\mu\text{M}$ | Autofluorescence of lipofuscin after treatment with bioactive substances | |
| --- | --- | --- | --- |
|  |  | Quercetin | Kaempferol |
| 1 | Control | $17613.0 \pm 185.9$ | |
| | 10 | $18183.0 \pm 133.1$ | $17948.0 \pm 179.4$ |
| | 50 | $17553.0 \pm 141.4$ | $17050.0 \pm 170.1$ |
| | 100 | $18037.0 \pm 179.6$ | $17728.0 \pm 165.4$ |
| | 200 | $17267.0 \pm 325.5$ | $16530.0 \pm 442.3$ |
| 5 | Control | $17801.0 \pm 264.0$ | |
| | 10 | $18664.0 \pm 518.6$ | $18986.0 \pm 629.9$ |
| | 50 | $17860.0 \pm 204.4$ | $16608.0 \pm 192.7$ |
| | 100 | $16279.0 \pm 524.9$ | $18295.0 \pm 791.0$ |
| | 200 | $16200.0 \pm 257.0$ | $15965.0 \pm 319.1$ |
| 10 | Control | $19274.0 \pm 175.5$ | |
| | 10 | $20418.0 \pm 793.8$ | $21217.0 \pm 567.8$ |
| | 50 | $19195.0 \pm 342.2$ | $18282.0 \pm 990.5$ |
| | 100 | $16671.0 \pm 582.5$ | $20241.0 \pm 471.2$ |
| | 200 | $16616.0 \pm 569.8$ | $16827.0 \pm 336.8$ |
| 15 | Control | $19662.0 \pm 126.5$ | |
| | 10 | $19284.0 \pm 443.9$ | $20048.0 \pm 340.7$ |
| | 50 | $18315.0 \pm 295.8$ | $17234.0 \pm 901.6$ |
| | 100 | $15716.0 \pm 554.0$ | $19170.0 \pm 547.1$ |
| | 200 | $15741.0 \pm 495.7$ | $15950.0 \pm 327.4$ |

| Nematode incubation time, days | Concentration of bioactive substance, $\mu\text{M}$ | Autofluorescence of lipofuscin after treatment with bioactive substances | |
| --- | --- | --- | --- |
|  |  | Quercetin | Kaempferol |
| 1 | Control | $17613.0 \pm 185.9$ | |
| | 10 | $18183.0 \pm 133.1$ | $17948.0 \pm 179.4$ |
| | 50 | $17553.0 \pm 141.4$ | $17050.0 \pm 170.1$ |
| | 100 | $18037.0 \pm 179.6$ | $17728.0 \pm 165.4$ |
| | 200 | $17267.0 \pm 325.5$ | $16530.0 \pm 442.3$ |
| 5 | Control | $17801.0 \pm 264.0$ | |
| | 10 | $18664.0 \pm 518.6$ | $18986.0 \pm 629.9$ |
| | 50 | $17860.0 \pm 204.4$ | $16608.0 \pm 192.7$ |
| | 100 | $16279.0 \pm 524.9$ | $18295.0 \pm 791.0$ |
| | 200 | $16200.0 \pm 257.0$ | $15965.0 \pm 319.1$ |
| 10 | Control | $19274.0 \pm 175.5$ | |
| | 10 | $20418.0 \pm 793.8$ | $21217.0 \pm 567.8$ |
| | 50 | $19195.0 \pm 342.2$ | $18282.0 \pm 990.5$ |
| | 100 | $16671.0 \pm 582.5$ | $20241.0 \pm 471.2$ |
| | 200 | $16616.0 \pm 569.8$ | $16827.0 \pm 336.8$ |
| 15 | Control | $19662.0 \pm 126.5$ | |
| | 10 | $19284.0 \pm 443.9$ | $20048.0 \pm 340.7$ |
| | 50 | $18315.0 \pm 295.8$ | $17234.0 \pm 901.6$ |
| | 100 | $15716.0 \pm 554.0$ | $19170.0 \pm 547.1$ |
| | 200 | $15741.0 \pm 495.7$ | $15950.0 \pm 327.4$ |

Based on the data in Table 1, we calculated the differences in fluorescence on each day of spectrometric measurements, subtracting the values obtained on the first day under the same conditions. Figures 1 and 2 show the effects of the tested bioactive substances on the accumulation of lipofuscin, depending on the time of nematode incubation (1–15 days).

**Figure 2.**
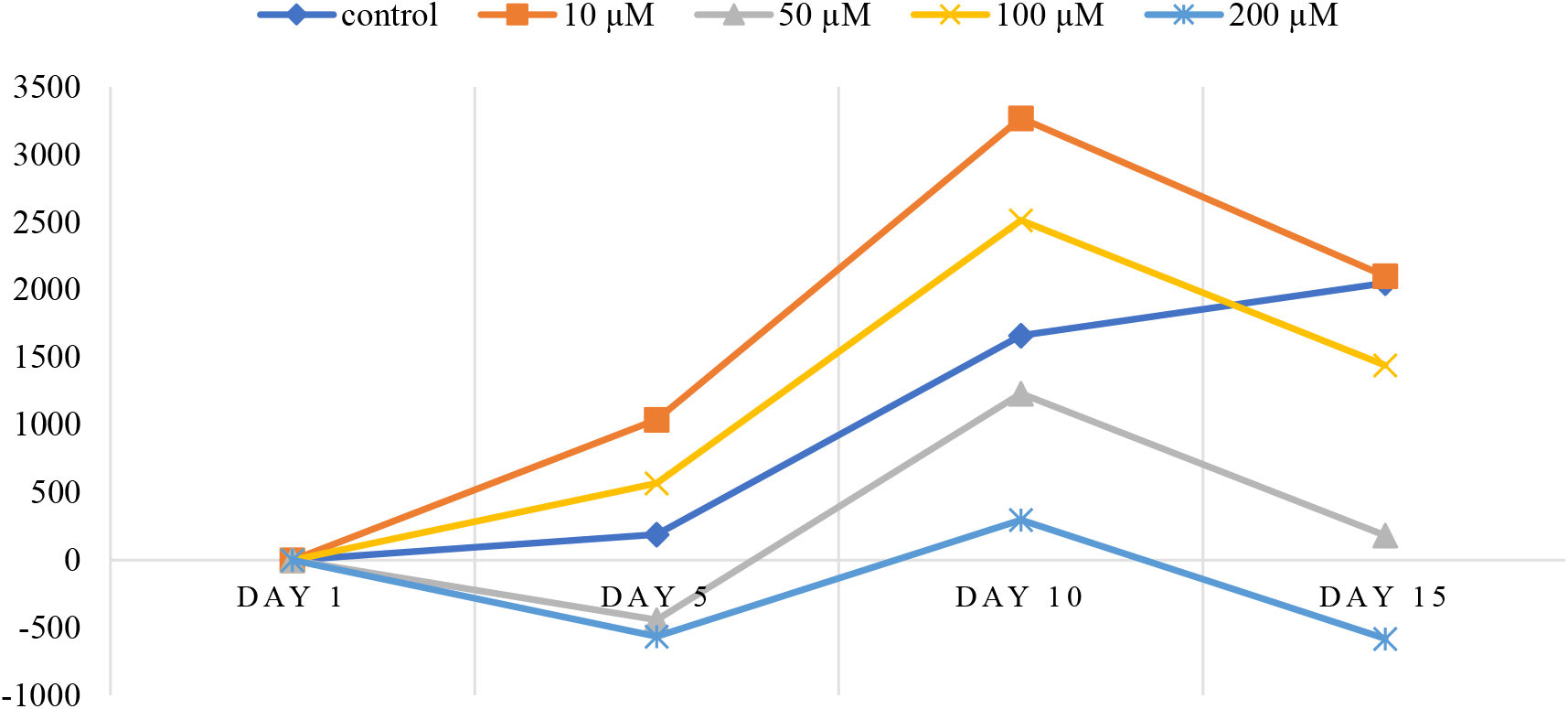
Accumulation of lipofuscin in *C. elegans* treated with kaempferol isolated from the *Ginkgo biloba* L. suspension culture extract

We found that quercetin and kaempferol effectively reduced lipofuscin accumulation in the nematodes on day 5 after the treatment. Both compounds steadily restrained the accumulation of lipofuscin until the end of the 15-day experiment. Quercetin at concentrations of 100 and 200 μM showed a significant decrease in the age-related pigment, compared to the control. Kaempferol at concentrations of 50 and 200 μM also reduced the level of lipofuscin. The 200 μM concentration of quercetin was 1.5 times more effective in reducing the accumulation of the age pigment over 15 days than the same concentration of kaempferol.

## Conclusion

We studied the effect of quercetin and kaempferol isolated from the *Ginkgo biloba* L. suspension culture extract on the accumulation of the age pigment lipofuscin in *C. elegans*. At a concentration of 200 μM, these bioactive substances showed an effective decrease in lipofuscin throughout the entire 15-day experiment, compared to the control. Quercetin was 1.5 times more effective in reducing lipofuscin levels than kaempferol. Thus, our study showed that these individual bioactive substances can inhibit lipofuscin, a factor that limits cell functionality during aging.

## Funding

*The study was part of the state assignment “Plant polyphenols of the Siberian Federal District: assessment of the molecular and spatial structure of substances, characterization of biofunctional properties and toxicological safety indicators on in vivo model systems” (project FZSR-2023-0002)*.

*The equipment was provided by the Center for Collective Use “Instrumental Methods of Analysis in Applied Biotechnology” at Kemerovo State University*.

## Notes

### Competing Interest Statement

The authors have declared no competing interest.

